# Gencube: Efficient retrieval, download, and unification of genomic data from leading biodiversity databases

**DOI:** 10.1101/2024.07.18.604168

**Authors:** Keun Hong Son, Je-Yoel Cho

## Abstract

**Motivation:** With the daily submission of numerous new genome assemblies, associated annotations, and experimental sequencing data to genome archives for various species, the volume of genomic data is growing at an unprecedented rate. Major genomic databases are establishing new hierarchical structures to manage this data influx. However, there is a significant need for tools that can efficiently access, download, and integrate genomic data from these diverse repositories, making it challenging for researchers to keep pace.

**Results:** We have developed *Gencube*, a command-line tool with two primary functions. First, it facilitates the utility of genome assemblies, related annotations, gene set sequences, and cross-species data from various leading biodiversity databases. Second, it helps researchers intuitively explore experimental sequencing data that meets their needs and consolidates the metadata of the retrieved outputs.

**Availability and implementation:** *Gencube* is a free and open-source tool, with its code available on GitHub: https://github.com/snu-cdrc/gencube.

## 1 Introduction

The transformative potential of comparative omics within and across species lies in its ability to uncover deep evolutionary patterns, illuminate genetic diversity, and drive genomic innovation, revolutionizing our understanding of complex biological systems. To achieve this, we need reference-quality genome assemblies for a wide range of species, along with comprehensive annotations, and a vast amount of experimental sequencing data using various techniques.

Technological breakthroughs, advanced computational methods, and decreasing sequencing costs have expanded the breadth and depth of genome generation efforts. The breadth of these initiatives is evident in ambitious biodiversity projects like the Earth BioGenome Project (EBP) and its affiliates, including the Vertebrate Genomes Project and Zoonomia (Lewin et al., 2018; Rhie et al., 2021; Zoonomia Consortium, 2020). The depth of these efforts is demonstrated by the enhanced understanding of genomic variation within species, as seen in the Human Pangenome Reference Consortium (HPRC) and Dog10K (Liao et al., 2023; Meadows et al., 2023).

Newly generated genomes have been submitted to GenBank, and traditional databases like RefSeq, UCSC Genome Browser, and Ensembl have been pivotal in providing curated genomes and annotations predicted by various approaches (Sayers et al., 2024; O’Leary et al., 2016; Nassar et al., 2023; Martin et al., 2023). However, to quickly provide access to the rapidly increasing number of genomes, new repositories like UCSC GenArk (Clawson et al., 2023) and Ensembl Rapid Release were launched. Around the time, Zoonomia released gene annotations inferred using the TOGA (Tool to infer Orthologs from Genome Alignments) method (Kirilenko et al., 2023). However, programmatic access to all these new repositories is not available. Additionally, these groups use different chromosome naming conventions, making data integration time-consuming and cumbersome for researchers. For these reasons, a program that provides comprehensive search across various databases, as well as capabilities for data download and integration, is critically needed to be useful to most life science researchers.

Currently, most experimental sequencing data is stored and managed by the International Nucleotide Sequence Database Collaboration (INSDC), which coordinates with SRA, ENA, and DDBJ (Arita et al., 2021). The key challenge lies in finding data suitable for specific research and obtaining the corresponding metadata. However, popular tools like *sra-tools, ffq*, and *fetchngs* are limited to accession inputs (NCBI SRA Toolkit Development Team, 2024; Gálvez-Merchán et al. 2023; Ewels et al., 2020). While programs like NCBI Entrez Direct and Entrez Programming Utilities (E-utilities) (Sayers et al., 2023) allow for keyword searches, they are not intuitive, require significant time to learn, and need programming knowledge to consolidate metadata from search results.

Here, we present *Gencube*, a Python-based, open-source command-line tool designed to streamline programmatic access to metadata and diverse types of genomic data from publicly accessible leading biodiversity repositories (Fig. 1). This software simplifies the retrieval, download, and standardization of genomic data, and facilitates the rapid exploration of experimental sequencing data, enabling researchers to efficiently collect datasets without resorting to labor-intensive and error-prone manual web methods. By enhancing data accessibility and reliability, *Gencube* empowers researchers to perform more accurate and effective omics data analyses.

**Figure 1.**
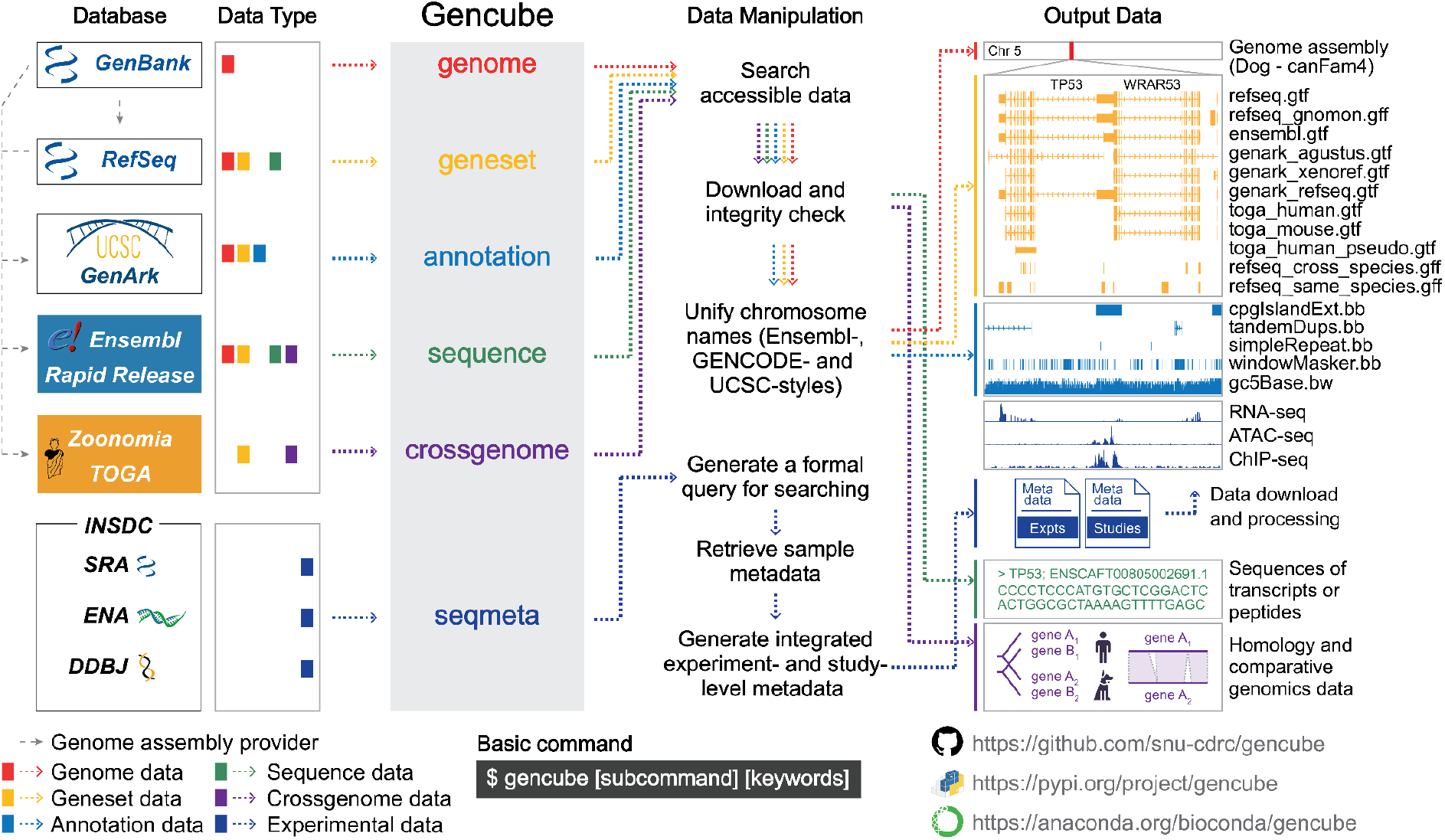
Overview of the *Gencube* workflow. The left panel lists the databases accessible through *Gencube*, along with the types of data retrieved. The middle panel illustrates the key subcommands and data manipulation processes. The right panel showcases examples of the output data that are downloaded and unified. The entire set of processes, from accessing the data to generating the final output, is color-coded according to the corresponding data type.

## 2 Description

Gencube consists of six key subcommands (Fig. 1), each dealing with different types of data:

- *gencube genome*: Fetches metadata and Fasta format files for genome assemblies.
- *gencube geneset*: Fetches GTF, GFF, or BED format files for gene annotations.
- *gencube annotation*: Fetches BigBed or BigWig format files for several types of genome annotations, such as gaps, GC percent, CpG islands, and repeats.
- *gencube sequence*: Fetches Fasta format files for transcripts or peptides sequences.
- *gencube crossgenome*: Fetches comparative genomics data, such as homology or codon- and protein-alignment of genes from different species.
- *gencube seqmeta*: Generates a formal search query for experimental sequencing data, retrieves the relevant metadata, and integrates it into experiment-level and study-level formats.

The first five collectively perform an initial search and fetch metadata of genome assemblies based on user requirements using NCBI E-utilities via Biopython (Cock et al., 2009). Researchers can use various forms of input accepted by NCBI Entrez, including scientific or common names of species, accessions, assembly names, UCSC names, such as homo_sapiens, human, GCF_029378435.1, GRCh38, and hg38. After this initial step, each subcommand accesses one or several public databases to check the availability of the targeted data for the searched genomes and displays the results in the terminal. Except for GenArk and TOGA, repositories provide MD5 checksum information to ensure data integrity. If the checksum of a previously downloaded or newly downloaded file differs from the server’s checksum, the download will be retried once to maintain data consistency.

Although the sequence information of genomes is the same across databases, the *genome* allows downloads from multiple sources because each database applies different masking methods. If masking affects the analysis, users can choose the appropriate source based on their needs and select between soft-masked, hard-masked, and unmasked genomes as required.

When analyzing data, it is crucial that the genome, geneset, and other annotations have consistent chromosome names. However, databases use various naming conventions, primarily falling into four categories: GenBank, RefSeq, Ensembl, and UCSC. The *gencube* standardizes chromosome names in files downloaded through the *genome, geneset*, and *annotation* according to user preferences, unifying them into Ensembl, GENCODE, or UCSC styles:

- Ensembl: Uses simple numeric and letter designations (e.g., 1, 2, X, MT). Unknowns use GenBank IDs.
- GENCODE: Uses “chr” prefix (e.g., chr1, chr2, chrX, chrM). Unknowns use GenBank IDs.
- UCSC: Also uses the “chr” prefix (e.g., chr1, chr2, chrX, chrM) but employs UCSC-specific IDs for unknowns, with limited use if UCSC IDs are not issued.

To modify the BigBed and BigWig files (binary format) downloaded by the *annotation*, they need to be converted to BED and BedGraph formats, respectively. After making the necessary modifications, these files are then restored to their original formats. In these processes, the UCSC genome browser utilities (*bigBedToBed, bigWigToBedGraph, bedToBigBed*, and *bedGraphToBigWig)* (Kent et al. 2010) are used instead of using a Python library.

The *seqmeta* also accesses sequencing data from the INSCD through NCBI Entrez. Initially, it generates a formal search query and implements all fields, properties, and filters available in Entrez as options (Fig. 2A). The input can include not only accession numbers but also a variety of relevant keywords. Additionally, wildcard (*) and the caret (^) at the end of the search terms can be used to include all entries containing the specified keyword (e.g., cancer* will include cancer, cancers, etc.) and perform exact word combination searches (e.g., liver_cancer^ will search for the exact phrase “liver cancer”).

**Figure 2.**
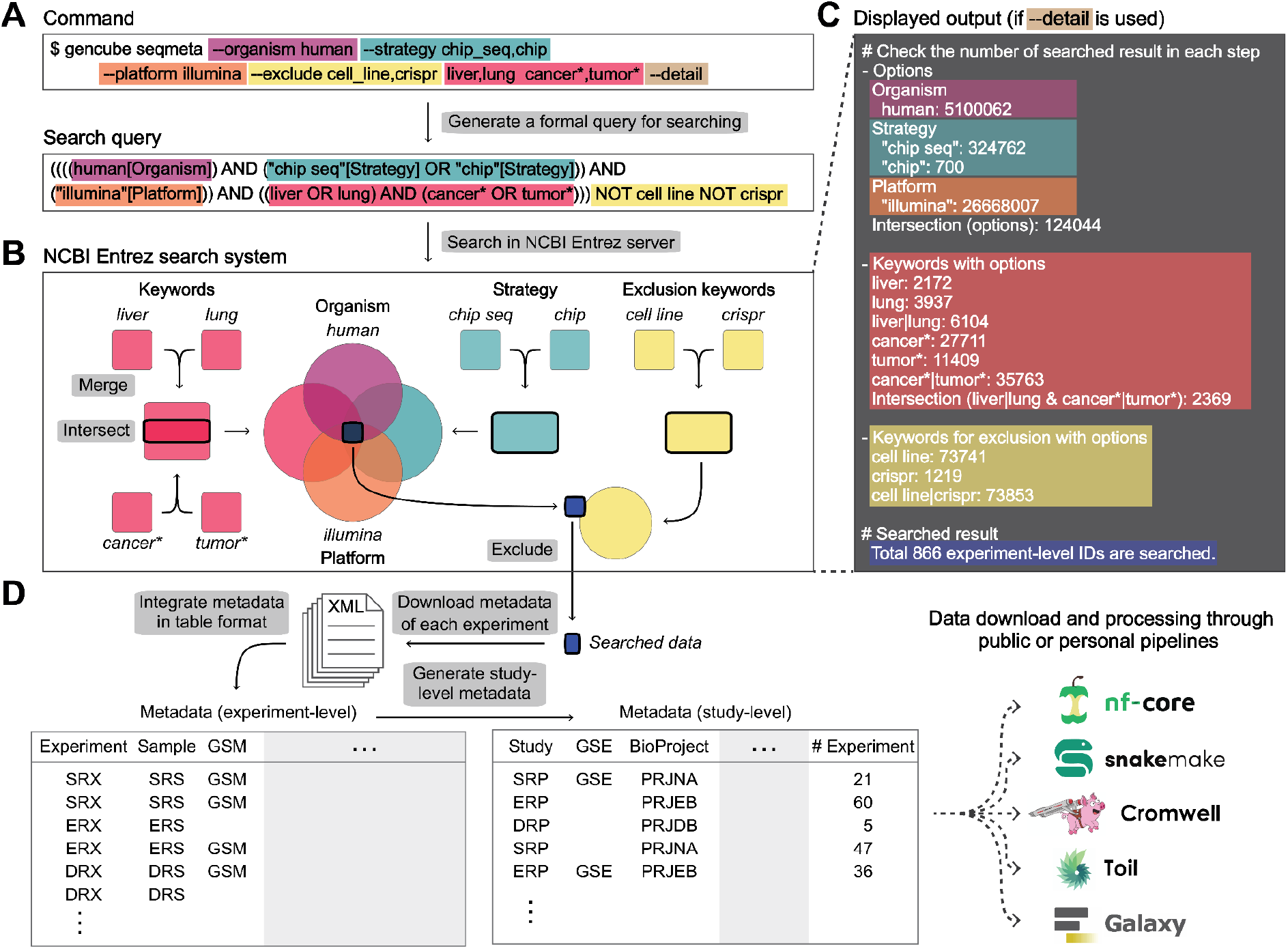
Schematic workflow of the *seqmeta* subcommand. (A) An example of the *seqmeta* command and its converted form as a search query. (B) Synopsis of data retrieval for the search query from NCBI Entrez. (C) Displayed output related to the data retrieval step. Each part of the command and the corresponding part in panels B and C are marked with matched colors. (D) The *seqmeta* downloads experiment metadata for all search results, extracts core information, integrates experiment-level output into a table format, and finally generates study-level output.

Depending on how the command is structured, Boolean operators (AND, OR, NOT) are used to either broaden or narrow the scope of the search, ultimately outputting the final set of experiment IDs (Fig. 2B). By displaying the number of results based on different options and keywords at each step in the terminal, it helps in making decisions about the query composition during the search process (Fig. 2C). Finally, for all the obtained IDs, metadata in XML format are collected and converted into table format for intuitive understanding by the user, creating both integrated experiment-level and study-level outputs (Fig. 2D).

*Gencube* utilizes well-established packages to ensure robust server interaction and data processing capabilities. Specifically, it employs *Pandas* (The Pandas Development Team 2023) for efficient data manipulation, *requests* (Reitz, 2024) and *BeautifulSoup4* (Richardson, 2024) for reliable web data access, and *tqdm* (da Costa-Luis et al., 2024) for enhanced result presentation. The software has undergone comprehensive testing on Linux/Unix and Mac OS (Darwin) platforms, confirming its cross-platform functionality and stability.

## 3 Usage and documentation

*Gencube* can be installed via the command line using ‘pip install gencube’. Alternatively, it can also be set up with conda. Users can invoke the help flag [-h] in the command line to receive detailed usage instructions. Additionally, the full manual, complete with examples, is available on the GitHub page at https://github.com/snu-cdrc/gencube. This ensures that users have access to thorough guidance for all subcommand functionalities.

## 4 Discussion

The number of genomes and annotations submitted annually from various consortia and projects is increasing exponentially. *Gencube* aims to consolidate these dispersed data sets to enable researchers to efficiently and quickly utilize them. Moreover, *Gencube* seeks to update biodiversity databases as they are created. While the most crucial information has been curated, the metadata submitted by researchers to INSDC is not yet perfectly standardized. Therefore, some manual selection of experiments from the obtained integrated metadata is necessary for research use.

In addition to INSDC, the Genome Sequence Archive (GSA) (CNCB-NGDC Members and Partners, 2023) is another major repository. Although smaller in scale compared to INSDC, GSA hosts a significant amount of sequencing data. Its web search logic is structured similarly to INSDC; however, programmatic search is not yet available. Once this feature is enabled, retrieval functionality for GSA will also be incorporated.

## Acknowledgments

We thank the EBP and its affiliates, GenArk, Ensembl Rapid Release, and science X (formerly known as Twitter), for their inspirational support. We extend our gratitude to Cookiecutter for providing the template for a Python package, which facilitated the swift and efficient development of this tool. We also thank the anonymous reviewers for their insightful comments and valuable suggestions.

## Funding

This work was supported by the Science Research Center (SRC) Program (grant no. NRF-2021R1A5A1033157 awarded to J.-Y.C.) under the Directorate for Basic Research in Science and Engineering, awarded as part of the Comparative Medicine Disease Research Center (CDRC) initiative, through the National Research Foundation (NRF) funded by the Korean government’s Ministry of Science and ICT.

## Data availability

All data and code relevant to this manuscript are available at: https://github.com/snu-cdrc/gencube

## Notes

https://github.com/snu-cdrc/gencube

### Competing Interest Statement

The authors have declared no competing interest.

